# Inhibitory Interneurons Modulate Neurovascular Coupling Primarily Through Indirect Circuit Mechanisms

**DOI:** 10.64898/2026.08.25.743501

**Authors:** Laurianne Zana, Milene R. Malheiros-Lima, Antoine Malescot, Éric Martineau, Ravi L. Rungta

## Abstract

Neurovascular coupling (NVC) links neuronal activity to local hemodynamics, underlies functional imaging signals such as fMRI, and is often disrupted in neurological disorders. Although inhibitory interneurons can directly signal to blood vessels, their relative contribution to NVC during sensory processing remains unclear. Here, we combined mesoscale cell-type-specific calcium imaging, hemodynamic imaging, and chemogenetic silencing to determine how parvalbumin-expressing (PV) and somatostatin-expressing (SOM) interneurons shape functional hyperemia in the mouse barrel cortex. During single-whisker stimulations, PV and SOM activity exhibited strong spatial co-variation with local hemodynamic responses across the barrel field. Silencing PV interneurons produced variable changes in local hemodynamic responses that closely tracked excitatory activity while disproportionately broadening the spatial spread of the hemodynamic response, whereas SOM silencing exerted comparatively modest effects. Together, these findings suggest that NVC predominantly reflects overall circuit activity, even when inhibitory signaling is broadly impaired.

## INTRODUCTION

Brain activity triggers an increase in local energy supply by signaling to blood vessels, a process known as neurovascular coupling (NVC). NVC is initiated on sub-second timescales through the detection and integration of neuronal activity across an interconnected capillary network^1^. These local signals backpropagate along the vasculature via waves of endothelial hyperpolarization^2,3,4^, driving dilation of upstream arterioles and proximal contractile capillary segments^1,2,3,4,5,6^. In addition to the propagation of signals across the capillary–arteriole network, vasoactive compounds released by neurons and glial cells directly regulate vascular tone^7,8,9^.

At the mesoscale, hemodynamic signals closely track bulk neuronal activity across the cortex^1,10^, exhibiting stronger correlations with local field potentials compared to spiking activity^11,12^. These observations have established the prevailing view that hemodynamic signals reflect local synaptic and circuit activity, underpinning the interpretation of functional imaging modalities such as fMRI. However, an emerging body of work has challenged this framework by proposing that inhibitory interneurons may be dominant regulators of functional hyperemia through direct vasoactive interactions with the cerebral vasculature^13–24^. If correct, this model would imply that hemodynamic signals may not primarily reflect overall neuronal activity, but instead be disproportionately influenced by the activity of specific inhibitory neuronal populations. Nonetheless, this emerging view heavily relies on studies employing direct optogenetic activation or other non-physiological stimulation paradigms. For example, direct activation of individual GABAergic interneurons *in vitro* is sufficient to induce either constriction or dilation of neighboring micro vessels^13^. Likewise, optogenetic activation of large interneuron populations i*n vivo* can elicit robust hemodynamic signals, despite a decrease in local excitatory neuronal activity^14,15^. Moreover, selective modulation of genetically defined interneuron subtypes can produce distinct vascular responses^16,17,18,19^, suggesting the existence of subtype-specific NVC pathways. While these studies have provided important mechanistic insights, they largely overlook the complex influence of interneurons on cortical circuit activity during physiological sensory processing, leaving unresolved whether interneurons regulate functional hyperemia through direct vascular signaling or indirectly through their effects on local network activity.

Here, we combined mesoscopic cell-type-specific imaging with chemogenetic silencing to determine how somatostatin (SOM) and parvalbumin-expressing (PV) interneurons (together comprising ∼70% of cortical interneurons) shape the spatiotemporal organization of neurovascular coupling to natural sensory stimulations. Using single-whisker deflections and optical imaging of the mouse barrel cortex, we show that sensory-evoked activity patterns from distinct interneuron subclasses can be reliably mapped at the mesoscale alongside local hemodynamic signals. PV and SOM activity exhibited strong co-variation with hemodynamic responses across the barrel field in both awake and sedated mice, with bulk Ca²⁺ signals showing a similar correspondence to hemodynamic signals as previously reported for excitatory neurons^1^. Silencing experiments further revealed distinct circuit contributions of interneuron subclasses to functional hyperemia. Whereas SOM inhibition produced comparatively modest effects on sensory-evoked neuronal and vascular responses, PV silencing altered the temporal evolution and spatial spread of the hemodynamic response in close correspondence with changes in excitatory activity. Together, these findings suggest that major inhibitory interneuron subclasses shape NVC predominantly through their regulation of local circuit dynamics rather than through dominant direct vascular control.

## RESULTS

### Mesoscopic imaging of PV and SOM interneuron activity and sensory-evoked hemodynamic responses

The somatotopic organization of the mouse primary whisker cortex serves as a powerful model for probing the spatial dynamics of neurovascular signals, as each whisker maps onto a spatially confined cortical unit, giving rise to the characteristic whisker “barrel” structure^25,26^ (**Fig. 1a**). We previously leveraged this organization to investigate the spatial relationship between excitatory Ca²⁺ signals and hemoglobin changes^1^, suggesting that a similar approach could be used to examine interneuron-NVC relationships. We therefore set out to examine the spatio-temporal relationship between interneuron subtype activity and hemoglobin signals at the mesoscale using wide-field optical imaging in *PV-Cre::GCaMP6f-floxed* and *SOM-Cre::GCaMP6f-floxed* mice implanted with chronic glass windows. Given the relatively sparse distribution of interneurons compared to excitatory neurons in the cortex, we first verified that bulk mesoscopic signals could be reliably detected with sufficient signal-to-noise to resolve barrel-specific responses. Using a three-LED optical imaging system, we simultaneously recorded interneuron bulk Ca²⁺ and hemodynamic responses, estimating total hemoglobin (HbT, ∼blood volume) from backscatter measurements of oxygenated (HbO) and de-oxygenated (HbR) hemoglobin, alongside GCaMP6f fluorescence in targeted interneuron cell populations (**Fig. 1a and Supplementary Fig. 1**). Brief stimulation (4 seconds, 5 Hz) of individual whiskers (E2 to B2) evoked rapid increases in both PV and SOM GCaMP6f signals, with spatially distinct activation areas corresponding to the stimulated whisker. Averaging across trials (∼15-30) revealed localized peaks in both SOM and PV Ca^2+^ signals within the corresponding associated barrel, enabling the automated alignment of a standardized barrel region of interest (ROI) mask to the centroid of each response (**Fig. 1b and Supplementary Fig. 1c**). These ROIs were then used to quantify and analyze the spread of the interneuron Ca^2+^ signals, as well as HbT changes across neighboring barrel columns in both dexmedetomidine-sedated and awake mice (**Fig. 1c, d).** Consistent with prior work^1,27,28^, hemodynamic responses were faster in awake compared to sedated mice, both in their onset and return to baseline. Together, these results demonstrate that interneuron subtype Ca^2+^ signals and hemodynamic responses can be reliably mapped at the mesoscale across the barrel cortex in both awake and sedated animals.

**Figure 1.**
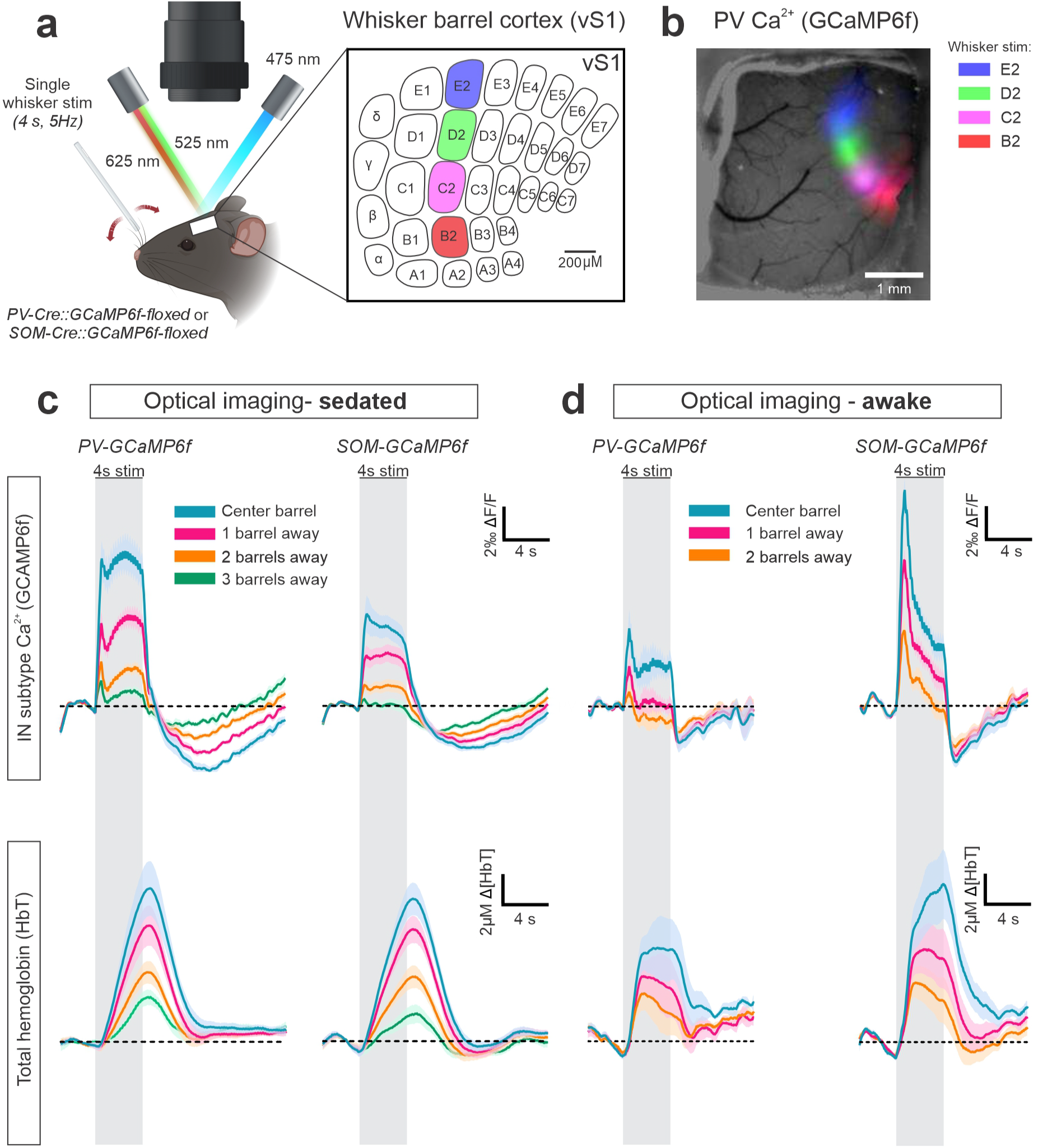
Mesoscale imaging of PV, SOM, and hemodynamic responses to single whisker deflections. **a,** Experimental setup schematic. Head-fixed *PV-Cre::GCaMP6f-floxed* and *SOM-Cre::GCaMP6f-floxed* mice were imaged with a wide-field optical imaging system. Four whiskers along the second column of the whisker barrel cortex (E2, D2, C2 and B2) were individually threaded into a glass capillary attached to piezoelectric element and stimulated at 5Hz for 4 seconds. Barrel map adapted from *Petersen, 2019*^25^. **b**, Color coded Ca^2+^ signals superimposed on image of craniotomy, showing average changes in GCaMP6f fluorescence in response to four individual whisker stimulations in a *PV::GCAMP6f* mouse. **c**, Average changes in GCaMP6f fluorescence (top) and HbT (∼blood volume, bottom) in *PV-GCaMP6f* (left, N=9, 3M and 6F) and *SOM-GCaMP6f* (right, N=12, 4M and 8F) dexmedetomidine-sedated mice, across the associated barrel ROI and its neighbors. **d**, Average changes in GCaMP6f fluorescence (top) and HbT (∼blood volume, bottom) in *PV-GCaMP6f* (left, N=4, 3M and 1F) and *SOM-GCaMP6f* (right, N=4, 2M and 2F) awake mice, across the associated barrel ROI and its neighbors. **c,d**, Data are represented as the mean ± Standard Error of the Mean (SEM). Shaded areas on traces represent the SEM. Grey bars represent the 4-second single whisker stimulation period. M = males, F = females.

### Spatial relationships between interneuron subtype activity and hemodynamic signals

We next examined the spatial relationship between interneuron-subtype activity and HbT signals across the barrel cortex. On average, in both awake and sedated mice, SOM and PV Ca^2+^ signals spread across neighboring barrel columns alongside HbT responses (**Fig. 1c, d**), reminiscent of our prior examination of excitatory-HbT relationships^1^. Examples of single whisker stimulations (E2) from individual mice show that while PV and SOM Ca^2+^ signals spread across multiple barrels, they often appeared more spatially confined than HbT signals (**Fig. 2a, b**), and the peak of the HbT elevation was occasionally located outside of the stimulated whisker barrel (e.g. **Fig. 2b),** likely due to the location of upstream the vasculature^1^. Analysis of the radial spread of normalized responses confirmed that, on average, both PV and SOM Ca²⁺ signals peaked within the barrel centroid and spread radially with spatial profiles closely resembling those of HbT signals in both sedated and awake animals (**Fig. 2c-f, Tables S1-S4**). However, PV Ca²⁺ responses were significantly more spatially confined than HbT signals in both conditions (distance to half-maximum), whereas the spread of SOM Ca²⁺ responses were comparable to that of HbT (**Fig. 2c-f, Tables S1-S4**).

**Figure 2.**
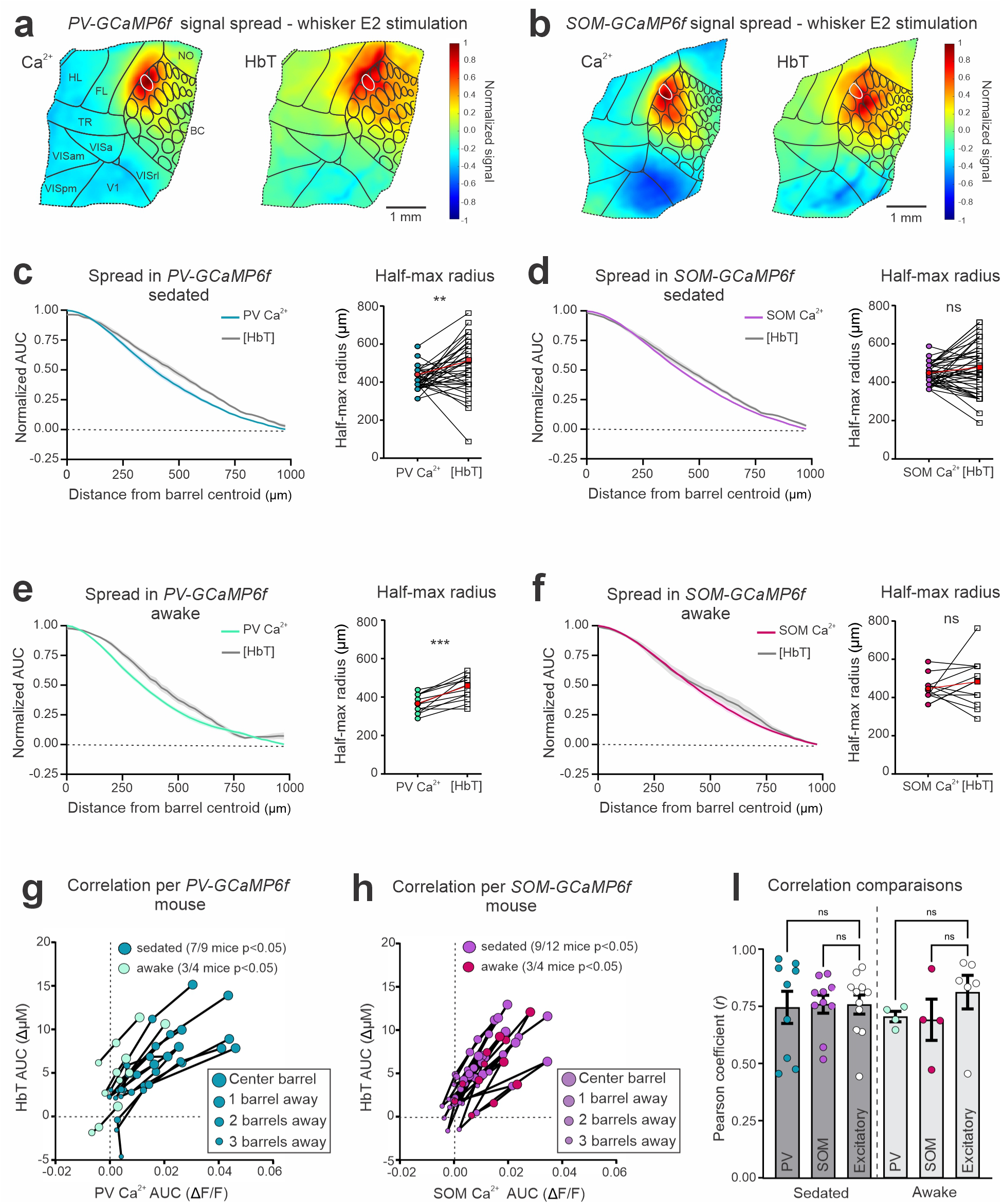
Spatial relationship between PV and SOM interneuron activity and hemodynamic signals. **a,b**, Example of Ca^2+^ (left) and hemoglobin (right) signals across the cortex in a *PV-GCaMP6f* (**a**) or *SOM-GCaMP6f* (**b**) sedated mouse, during the E2 whisker stimulation. **c-f**, Radial spread of interneuron Ca^2+^ and HbT responses (AUC) from the stimulated barrel centroid, and values of the half-max radius (μm) from both sedated and awake *PV-GCaMP6f* (**c** and **e** respectively) and *SOM-GCaMP6f* (**d** and **f** respectively) mice. Red datapoints represent the group average **g,h**, Changes (AUC) in Ca^2+^ (ΔF/F) and HbT (µM) averaged per barrel distance for each *PV-GCaMP6f* (**g**) and *SOM-GCaMP6f* (**h**) mouse, sedated and awake. Ratios indicate the number of mice for which the correlation between the Ca^2+^ and HbT signals across all barrel ROIs had a statistical significance of *p*<0.05, by two-tailed Pearson’s correlation. For AUC changes, AUC _t=4-8s_ was used for Ca^2+^ signals and AUC _t=4-10s_ was used for HbT. **i**, Comparison of the Pearson’s correlation coefficient (*r*) for individual animals between changes in HbT and Ca^2+^ for different cell types, in either sedated or awake conditions. Excitatory neuron data from *Thy1-jRGECO1a* mice and previously published in Martineau et al.^1^ * *p*<0.05 by one-way ANOVA with Sidak’s multiple comparison test. Data are represented as the mean ± SEM.

Despite these subtle differences in spread, the spatial distribution of Ca²⁺ and HbT responses across whisker barrels was highly correlated within individual animals, both awake and sedated, with significant relationships observed in the majority of mice for both PV (7/9 for sedated and 3/4 for awake) and SOM (9/12 for sedated and 3/4 for awake) interneurons (**Fig. 2g, h)**. The corresponding Pearson correlation coefficients (r) were not significantly different from those previously observed for excitatory neurons^1^ (**Fig. 2i, Table S5**). Consistent with these findings, regression analyses that accounted for inter-animal variability revealed that PV, SOM, and excitatory neuron Ca^2+^ each explained a similar proportion of the variance in sensory-evoked HbT responses, both in awake and sedated mice (R^2^_adj_ = 0.56-0.69 for all cell types, **Supplementary Fig. 2e-j, Tables S6-S9**). Together, these results demonstrate that the spatial organization of PV and SOM interneuron activity closely mirrors sensory-evoked hemodynamic responses across the barrel cortex, comparable to excitatory neurons^1^, highlighting the need for cell type-specific manipulations to disentangle their respective contributions to NVC.

### Modulating PV and SOM interneuron activity using a chemogenetic strategy

To causally determine the contribution of distinct interneuron subtypes to NVC, we next attempted to chemogenetically modulate PV and SOM activity using inhibitory Gi-DREADDs while simultaneously recording both excitatory and hemodynamic signals. Simultaneous imaging of excitatory activity was critical, as interneuron manipulation inevitably alters cortical network activity, making it otherwise impossible to determine whether changes in functional hyperemia arise from direct vascular actions of interneurons or indirectly through changes in excitatory drive. This strategy therefore allowed us to assess how interneuron silencing impacts NVC while accounting for changes in excitatory activity. *Thy1-jRGECO1a* mice, crossed to either *PV-Cre* or *SOM-Cre* animals, received a local injection of the Cre-dependent virus *AAV2/8-hSyn-DIO-hM4Di-IRES-mCitrine* on the day of the craniotomy (**Fig. 3a**). Three weeks following the surgery, the barrel cortex of each mouse was functionally mapped using single whisker stimulations, and the mouse was subsequently imaged with two-photon microscopy to confirm and locate the area of hM4Di expression via mCitrine fluorescence. This step served to validate the presence of hM4Di-expression in the barrel whose associated whisker would be stimulated in all future imaging experiments (**Fig. 3a**). We first validated our ability to selectively modulate PV and SOM interneurons with the DREADD agonist deschloroclozapine (DCZ, 100 μg/kg, IP), by measuring changes in the activity of L2/3 pyramidal cell spontaneous activity in sedated animals (**Fig. 3b**). DCZ was chosen over other DREADD agonists due to its reported higher selectivity^29,30^, and adverse interactions, in our hands, between CNO and dexmedetomidine sedation. DCZ administration in PV-hM4Di-expressing mice, triggered an increase in excitatory neuronal activity, marked by an elevation of spontaneous Ca^2+^ transient frequency within the soma of L2/3 Thy-1 neurons (**Fig. 3c, d, Tables S10**). In contrast, SOM interneuron silencing led to a small but significant decrease in L2/3 excitatory neuron somatic transients 20 minutes post-DCZ injection (**Fig. 3e, Table S11**), an effect that may appear counterintuitive but is consistent with the disinhibitory influences via reduced SOM–PV interactions^31^ and the preferential targeting of dendrites by SOM interneurons^32,33^, which may have a weaker impact on somatic Ca^2+^ compared to the somatic inhibition exerted by PV interneurons. Control vehicle injections did not have any effect on spontaneous excitatory Ca^2+^ transients (**Supplementary Fig. 3, Tables S26,27**). These results highlight the differing effects of SOM and PV interneuron silencing on spontaneous excitatory neuronal activity, reflecting the distinct connectivity profiles of each interneuron subtype within local circuitry.

**Figure 3.**
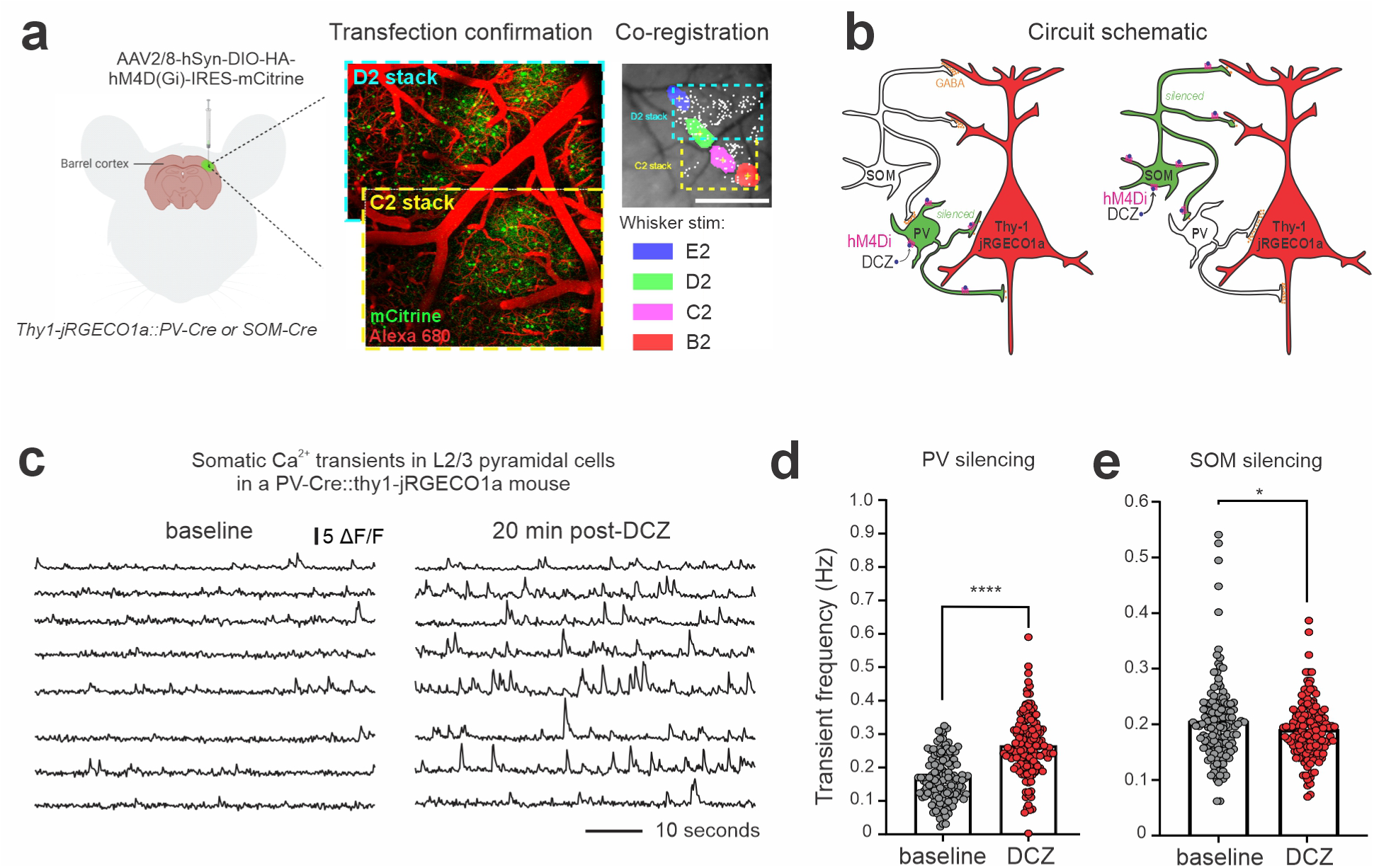
Targeting barrel cortex interneuron subtypes with Gi-DREADDS. **a**, Silencing experiment schematic. The Cre-dependent *AAV2/8-hSyn-DIO-HA-hM4D(Gi)-IRES-mCitrine* was locally injected in the barrel cortex of *PV-Cre::thy1-jRGECO1a* or *SOM-Cre::thy1-jRGECO1a* mice. Viral expression was confirmed via localization of mCitrine-positive interneurons using two-photon microscopy. Co-registration of the vasculature was used to align the area of mCitrine expression onto the barrel map. **b**, Schematic of the circuit. PV and SOM interneurons, are transduced with the hM4D(Gi) receptor and co-express mCitrine (green). To selectively silence PV or SOM interneurons, DCZ (100 μg/kg) is intraperitoneally injected (IP) to activate the hM4D(Gi) receptor while recording the impact on excitatory neuron Ca^2+^ signals, via jRGECO1a fluorescence (red). **c**, Example of spontaneous Ca^2+^ transients measured in the somas of eight L2/3 pyramidal cells before (left) and 20 minutes post-DCZ injection (right) in an hM4Di-expressing *PV-Cre::thy1-jRGECO1a* mouse, measured in ΔF/F. **d,e**, Ca^2+^ transient frequency in Hz, before and after DCZ injection in hM4Di-expressing *PV-Cre::thy1-jRGECO1a* (**d**, 6 mice, N=6, 3M and 3F) and *SOM-Cre::thy1-jRGECOa1* (**e**, N=6, 6M) sedated mice.\**p*<0.05 and \*\*\*\**p*<0.0001 by two-tailed paired t-tests. Between 48 and 143 somas per mouse.

### PV interneuron silencing modulates hemodynamic responses indirectly through changes in excitatory circuit activity

We next sought to determine how these two broad populations of interneurons shape the hemodynamic response during sensory processing. Using wide-field imaging, we first tested how PV interneuron silencing altered the dynamics of NVC, by measuring mesoscale excitatory neuronal activity and hemoglobin changes in response to 4 second whisker stimulations before and 20 minutes after DCZ administration. Neither vehicle (saline) injections in hM4Di-expressing mice, nor DCZ injections in hM4Di-negative mice, affected neuronal activity or the positive phase of the hemodynamic response (**Supplementary Fig. 4a-d, Tables S28-33**). Surprisingly though, I.P injection of DCZ, or vehicle alone, evoked an increase in the post stimulus undershoot phase of the hemodynamic response of sedated mice (**Supplementary Fig. 4b-d**). Therefore, the following data was only analyzed during the positive phase of the hemodynamic response (0 - 6 seconds following the onset of stimulation). Both the excitatory neuronal Ca^2+^ and HbT responses appeared to increase in the early phase of the response following DCZ injection in hM4Di-expressing *PV-Cre::thy1-jRGECO1a* mice (**Fig. 4a**). However, we observed substantial variability across mice, as illustrated by three individual examples (**Fig. 4b**). Notably, changes in excitatory activity systematically appeared to correspond with changes in the dynamics of the HbT signal, particularly in the early phase: mice showing larger and faster excitatory responses also exhibited stronger and faster HbT signals (left two examples), whereas animals with minimal excitatory change showed no HbT effect (single example shown on right) (**Fig. 4b**). The observed variability was linked to differences in viral expression spread across barrel columns following the intracortical AAV injection. In mice where hM4Di expression was mostly confined within one-barrel (i.e. low spread expression), localized PV silencing had no impact on either the neuronal activity or the hemodynamic response (**Fig. 4c-e, Tables S12,13**). Conversely, in mice where hM4Di expression spread through at least 2 barrels (i.e., widespread expression; **Fig. 4f-h, Tables S14,15**), PV silencing resulted in an increase in the early phase of the neuronal response (t=4-4.5 seconds), which was matched by a larger increase in blood volume (t=5.3-6.3 seconds). Importantly, successful DCZ-mediated PV suppression was confirmed in mice from both low spread and high spread expression groups via measurements of increased spontaneous activity in the stimulated barrel (**Fig. 3**). Together, these results demonstrate that, at the mesoscale, the early hemodynamic response is tightly coupled to excitatory activity, with PV silencing exerting little direct influence beyond its effects on circuit dynamics.

**Figure 4.**
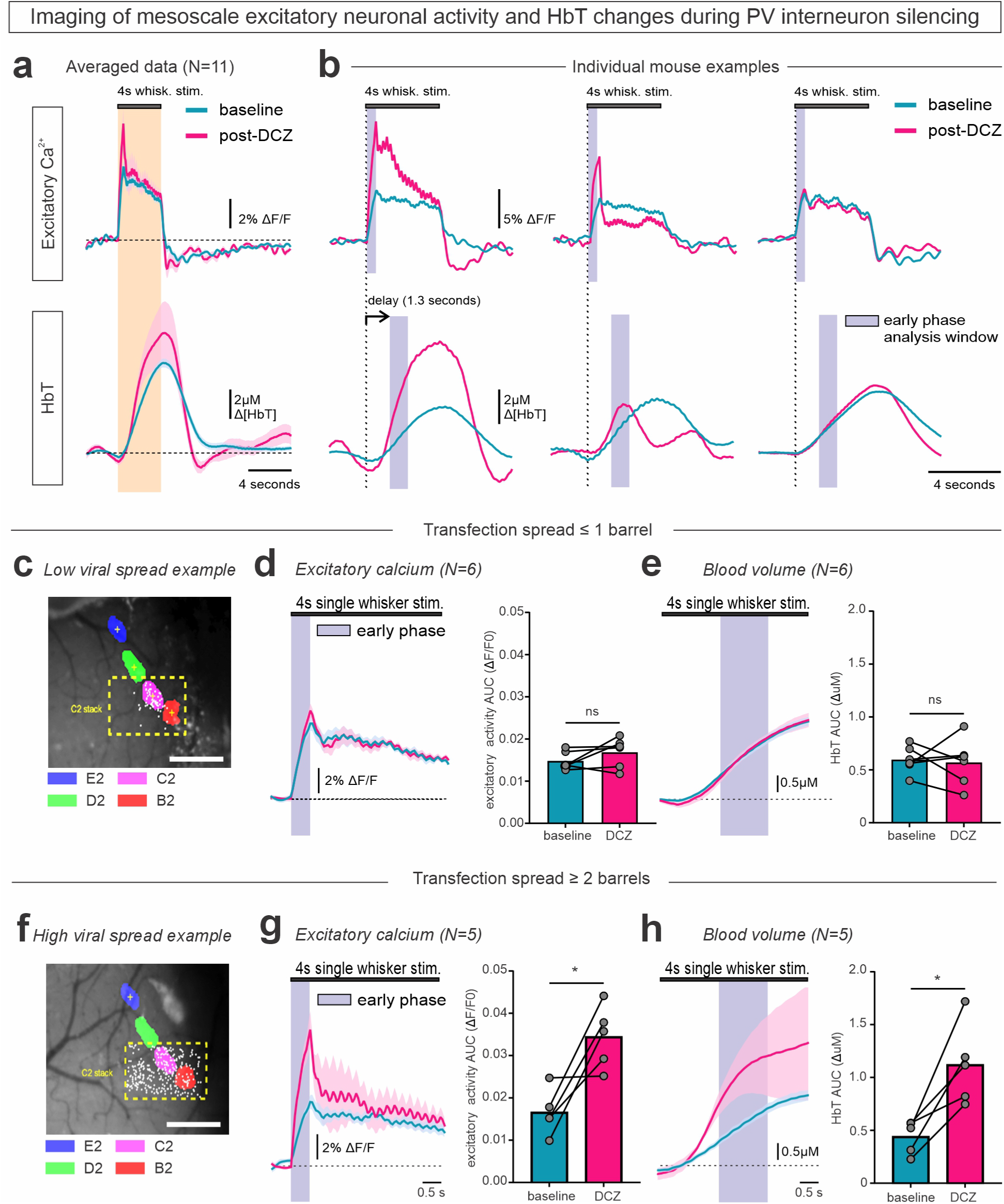
NVC tracks overall circuit dynamics following PV chemogenetic silencing. **a**, Average excitatory Ca^2+^ (top) and hemodynamic (bottom) response to single whisker stimulations before and 20 minutes post-DCZ injection in sedated hM4Di-expressing *PV-Cre::thy1-jRGECO1a* (N=11, 6M and 5F). **b**, Individual examples of excitatory Ca^2+^ (top) and hemodynamic (bottom) response to DCZ administration from three hM4Di-expressing *PV-Cre::thy1-jRGECO1a* mice. **c,** Example of a local expression pattern captured with two-photon microscopy and registered onto the S1 map of the same animal. **d,e,** Analysis of the early phase of the evoked excitatory Ca^2+^ response (**d**, AUC _t=4-4.5s_) and the hemodynamic response (**e**, AUC _t=5.3-6.3s_), in mice that exhibited low spread hM4Di expression (≤ 1 barrel away from the stimulated barrel, N=6, 2M and 4F). **f,** Example of a widespread expression pattern captured with two-photon microscopy and registered onto the S1 map of the same animal. **g,h**, Analysis of the early phase of the evoked excitatory Ca^2+^ response (**g**, AUC t=_4-4.5s_) and the hemodynamic response (**h**, AUC t=_5.3-6.3s_), in mice that exhibited widespread hM4Di expression (≥ 2 barrels away from the stimulated barrel, N=5, 4M and 1F). **c and f**, Dashed yellow box shows field of view of acquired two-photon z-stack. White dots indicate location of mCitrine positive neurons identified with two-photon imaging. **d,e,g,h**, * *p*<0.05 by two-tailed paired t-tests, N=6 for each group.

### PV chemogenetic silencing alters both the temporal and the spatial dynamics of NVC

Because PV interneurons provide rapid feedforward inhibition and contribute to the spatial sharpening of cortical activity^34,35^, we next asked whether PV silencing altered the temporal evolution and spatial spread of sensory-evoked neuronal and hemodynamic responses. First, the rise time of the Ca^2+^ and HbT responses were estimated by comparing the slope of the signals before and after PV silencing. In the high viral spread group, we observed a trend towards faster Ca^2+^ (**Fig. 5a, Table S16**) and HbT (**Fig. 5b, Table S17**) responses following DCZ injection. To determine if PV silencing altered the spatial relationship between excitatory Ca^2+^ and hemodynamic signals across space, donut-shaped ROIs were created by expanding circular masks around an initial ROI centered on the barrel centroid Signal AUC was then calculated within each donut before and after hM4Di receptor activation. Interestingly, in mice with widespread hM4Di expression, we observed that, while Ca^2+^ signals spread similarly across the barrel field following PV silencing (**Fig. 5c, d, Tables S18**), hemodynamic signals spread to a greater extent across S1 (**Fig. 5e, f, Tables S19**). However, we found no significant difference between the spatial spread of either signal pre and post-DCZ injection in the low hM4Di expression group, **(Supplementary Fig. 5, Tables S34,35)**. Overall, these results show that broad PV silencing enhances whisker-evoked excitatory activity, which correspondingly accelerates the onset of the hemodynamic response and increases its spatial spread across S1. These findings are consistent with stronger and faster neuronal responses recruiting broader increases in blood volume across the vascular network of the cortex.

**Figure 5.**
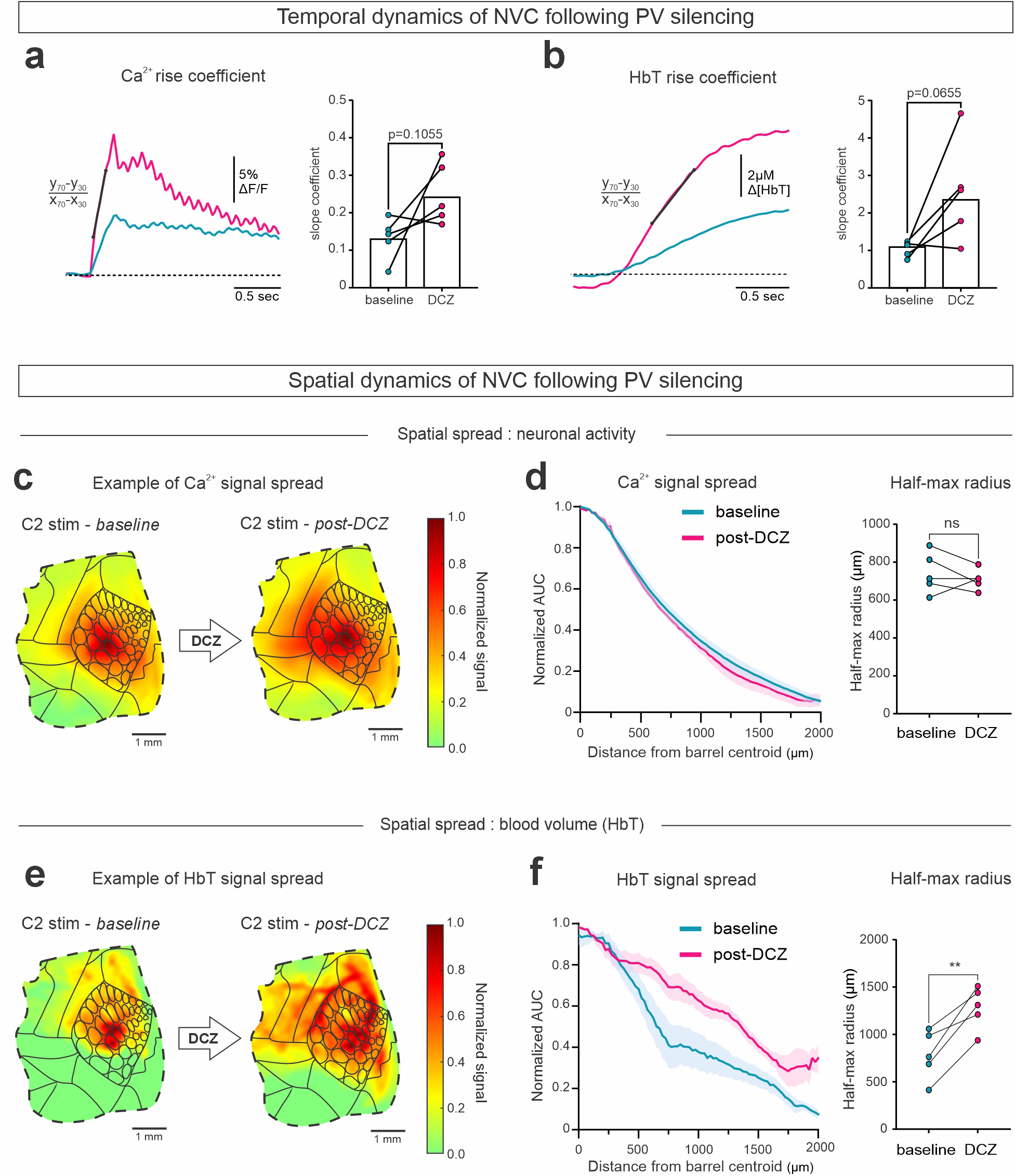
Spatiotemporal dynamics of NVC following PV chemogenetic silencing. **a,b,** Example traces from one mouse (left, widespread hM4Di-expression) and comparison of the slope coefficients (right) of the excitatory Ca^2+^ (**a**) and hemodynamic response (**b**) in sedated hM4Di-expressing *PV-Cre::thy1-jRGECO1a* mice before and after DCZ injection. **c,e**, Heatmaps represent an example of signal spread for excitatory Ca^2+^ (**c**) and HbT (**e**) in one hM4Di-expressing *PV-Cre::thy1-jRGECO1a* mouse, during a 4-second C2 whisker stimulation. Signals were normalized to the peak. **d,f.** Excitatory Ca^2+^ and HbT signal spread (AUC) from the whisker associated barrel centroid as well as the half-max radius (μm) in the hM4Di-expressing *PV-Cre::thy1-jRGECO1a* mice (wide-expression group). **a-f,** Data from the widespread hM4Di viral expressing group (N=5, 4M and 1F). **c-f**, All analysis was done on the early phase Ca^2+^ AUC _t=4-4.5s_ and hemodynamic response AUC _t=5.3-6.3s_.

### Chemogenetic silencing of SOM exerts minimal effects on NVC

Finally, we sought to determine the impact of SOM interneurons silencing on NVC during sensory processing, using the same chemogenetic approach as for PV cells. (**Supplementary Fig. 6a, b, Supplementary Tables S34,35**). On average, SOM silencing had no significant effect on either excitatory calcium or hemodynamic signals in sedated mice (**Fig. 6a, b, Tables S20,21**). As our mesoscale imaging experiments previously showed that SOM Ca^2+^ signals were larger in awake animals (**Fig. 1c, d**), it’s possible that SOM interneuron activity was not sufficient to reveal their effects under dexmedetomidine-sedation. Therefore, we further evaluated the effects of SOM silencing during single whisker stimulation in awake mice. Once again, SOM silencing had no marked impact on either excitatory Ca^2+^ or hemodynamic responses (**Fig. 6c, d, Tables S22,23**). Finally, to recruit a broader population of SOM interneurons, we examined NVC during simultaneous deflection of multiple whiskers (E2, D2, C2 and B2) in awake animals, which likewise revealed no significant effects of SOM silencing, but did show a trend towards increased neuronal and hemodynamic responses following DCZ injection in 3 out of 4 animals (**Fig. 6e, f, Tables S24,25**), consistent, if anything, with SOM inhibition influencing NVC predominantly through modulation of overall circuit activity. Similarly, one sedated animal from the widespread hM4Di viral group exhibited a notable increase in both neuronal and hemodynamic responses following DCZ injection compared to saline vehicle injection (**Supplementary Fig. 6c, d**). Taken together, these findings suggest that on average broad chemogenetic silencing of SOM interneurons exerts limited effects on mesoscale hemodynamic responses during sensory processing. While in some animals SOM silencing produced discernible changes, alterations in HbT paralleled changes in excitatory activity, consistent with modulation of NVC through changes in overall circuit activity.

**Figure 6.**
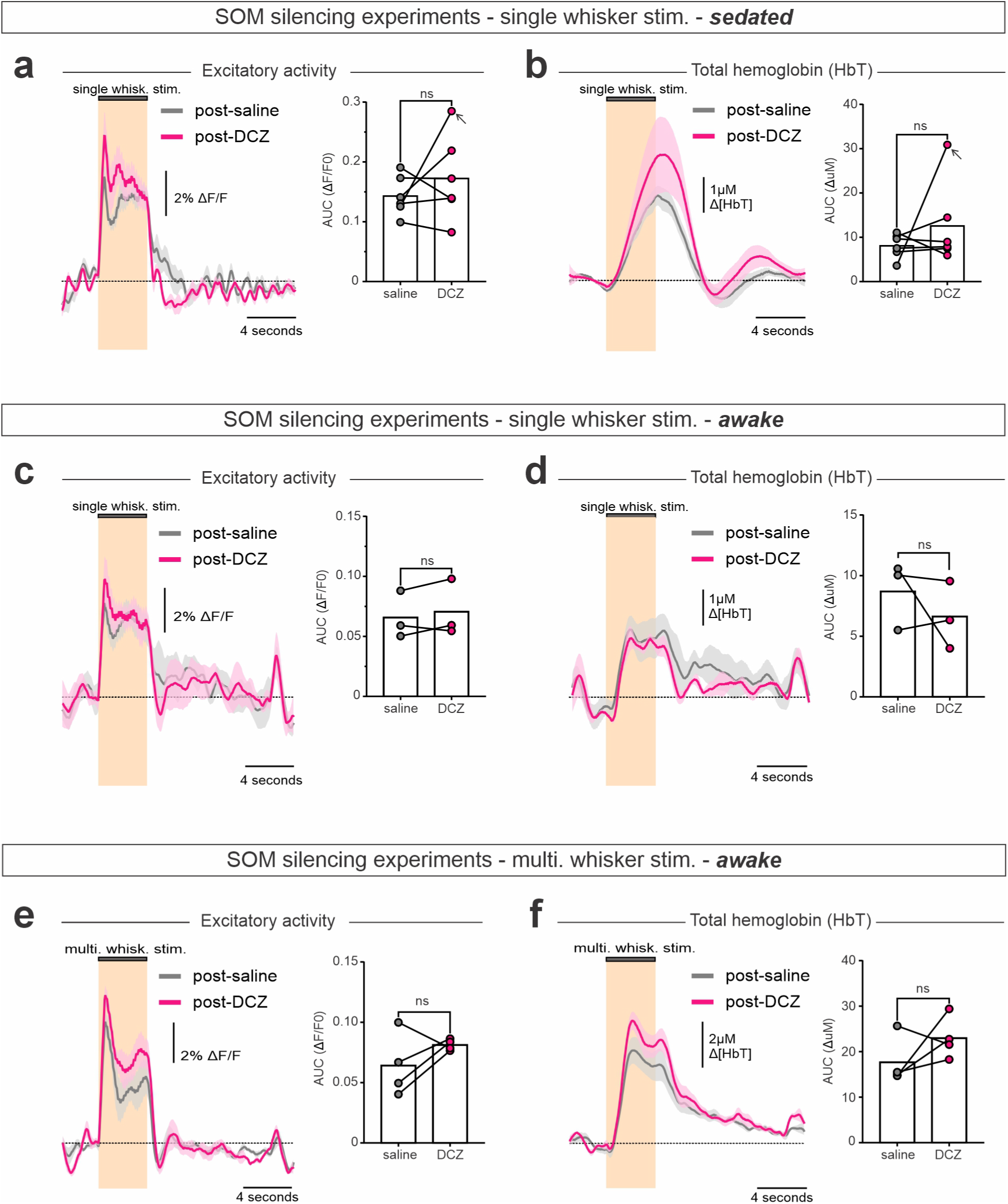
SOM chemogenetic silencing has minimal effects on NVC. **a,b,**) Average excitatory Ca^2+^ (**a**) and hemodynamic response (**b**) traces and AUCs in response to a single whisker stimulation after saline (vehicle) or DCZ injection in sedated hM4Di-expressing *SOM-Cre::thy1jRGECO1a* mice (N=6, 3M and 3F). The arrow indicates the mouse with the largest change in excitatory Ca^2+^ post-DCZ, in which the HbT response was also elevated. **c,d,**) Average excitatory Ca^2+^ (**c**) and hemodynamic response (**d**) traces and AUCs in response to a single whisker stimulation after vehicle or DCZ injection in awake hM4Di-expressing *SOM-Cre::thy1jRGECO1a* mice (N=3, 2M and 1F). **e,f,** Average excitatory Ca^2+^ (**e**) and hemodynamic response (**f**) traces and AUCs in response to the combined stimulation of a row of multiple whiskers (E2, D2, C2 and B2) after vehicle or DCZ injection in awake hM4Di-expressing *SOM-Cre::thy1jRGECO1a mice* (N=4, 4F). **a-f,** Data are represented as the mean ± the SEM. Shaded areas on signal traces represent the SEM. ns: *p*>0.05 by two-tailed paired t-test.

## DISCUSSION

In this study, we combined mesoscopic cell-type-specific imaging with chemogenetic silencing to determine how the two predominant inhibitory interneuron types contribute to NVC during a natural sensory stimulation. One notable finding of this study is that mesoscale activity patterns from distinct interneuron subclasses can be reliably resolved during single-whisker stimulation, something that was not trivial given their comparatively sparse representation in the cortex. Like excitatory neurons^1^, both PV and SOM activity exhibited spatial patterns that closely tracked local hemodynamic responses across the barrel field, in both awake and sedated animals. Together, these findings suggest that the close spatial relationship between interneuron subtype activity and hemodynamic responses reflects the integrated activity of the cortical circuit as a whole, rather than providing evidence for direct neurovascular signaling by specific cellular populations. By simultaneously imaging bulk excitatory neuronal activity and hemodynamic responses during chemogenetic silencing of specific interneuron subtypes, we were able to interpret changes in functional hyperemia in the context of accompanying changes in cortical network activity.

PV interneurons have been reported to evoke both vasoconstriction and/or vasodilation when optogenetically activated^16,19,22,23^, yet recent evidence suggests that silencing their activity has minimal impact on sensory evoked hemodynamic signals^24^, leaving their roles in NVC hard to interpret. Here, PV silencing revealed that their contribution to sensory-evoked NVC is mediated predominantly through their regulation of cortical circuit activity rather than through direct vascular signaling. Across animals, PV silencing produced variable effects, with changes in functional hyperemia closely paralleling accompanying changes in excitatory Ca^2+^ responses. Whereas widespread PV silencing markedly increased both sensory-evoked excitatory Ca^2+^ and blood volume responses, more spatially restricted silencing had little effect on either despite clear evidence of successful DREADD-mediated suppression of PV interneuron activity. The faster neuronal and hemodynamic responses following PV silencing is consistent with the well-established role of PV interneurons in feedforward inhibition within the barrel cortex^26^. Interestingly, PV silencing also broadened the spatial spread of functional hyperemia beyond the accompanying neuronal activity changes. One possible interpretation is that enhanced local excitatory activity more effectively engages retrograde vascular signaling mechanisms within the cerebrovascular network, allowing their amplification through endothelial signaling to propagate further upstream and across neighboring vascular territories^1,2,3,4,36,37^.

SOM interneurons have been strongly implicated in vascular regulation as subsets of these cells express vasoactive mediators such as nitric oxide (NO) and neuropeptides such as somatostatin and NPY ^13,15,16,17,18^. NO is a potent-vasodilator that is believed to be heavily implicated in NVC mechanisms^38^, and as such optogenetic or chemogenetic activation of nNOS neurons has been shown to evoke robust vasodilation^16,17,18,21,39^. In particular, a sparse subpopulation of SOM cells, labeled type-1 nNOS interneurons (∼0.5-2% of all GABAergic neurons in the cortex) have been suggested to exert disproportionately large effects on neurovascular responses during sensory stimulation^39^. However, new findings challenge the involvement of nNOS neurons in sensory-evoked NVC, with a recent study reporting no impact of NOS inhibition on functional hyperemia^21^. More importantly, local ablation of these type-1 nNOS neurons has only subtle effects on functional hyperemia which importantly were paralleled by decreases in bulk excitatory Ca^2+^ elevations^40^. Together, reinforcing our view of functional hyperemia primarily following overall circuit dynamics rather than the activity of a specific cell population.

Our study examined neurovascular coupling during physiologically relevant, medium-duration (4 second) whisker stimulations, a regime that primarily captures the initiation and early maintenance of sensory-evoked functional hyperemia. Within this temporal window, we found no evidence that broad chemogenetic inhibition of SOM interneurons substantially alters hemodynamic responses beyond their modest effects on local excitatory circuit activity. However, this does not exclude a role for specific SOM interneuron subpopulations under different conditions or timescales. Indeed, emerging evidence suggests that during prolonged stimulation lasting tens of seconds, somatostatin release can activate astrocytic signaling pathways that prolong the hemodynamic response^20^, consistent with studies implicating astrocytes in the slower components of neurovascular coupling^41,42^. Determining how this SOM-astrocyte pathway interacts with neuronal circuit dynamics, and whether it represents a direct vascular signaling axis or a secondary consequence of circuit alterations, will require further investigation.

Collectively, our findings refine current models of interneuron contributions to NVC. Rather than acting primarily through dominant direct vascular control, major inhibitory interneuron subclasses appear to shape hemodynamic responses largely through their influence on local circuit activity. Generally, these results highlight the importance of circuit-mediated effects in shaping alterations in NVC across conditions in which inhibitory signaling is disrupted, with important implications for the interpretation of functional imaging signals and for understanding how neurovascular signaling is altered in disease states characterized by inhibitory circuit dysfunction.

## METHODS

### Animals and ethic statements

All animal care and experiments were performed in accordance with the guidelines of the Canadian Council of Animal Care and the Comité de Déontologie sur l’Expérimentation Animale (CDEA) of the Université de Montréal. Male and female mice between 2 months and 8 months of age and weighing between 20 and 40 grams were used for this study. *PV-Cre(Cre/wt)::GCaMP6f(fl/wt)* and *SOM-Cre(Cre/wt)::GCaMP6f(fl/wt)* mice were obtained by crossing *Pvalb-IRES-Cre* (Jackson Labs, JAX #017320) and *SOM-IRES-Cre* (Jackson Labs, JAX #013044) mice with the Ai95 Cre-dependent *floxed-GCaMP6f* mouse line (Jackson Labs, JAX #028865). *PV-Cre(Cre/wt)::thy1-jRGECO1a(Tg/-)* and *SOM-Cre (Cre/wt)::thy1-jRGECO1a(Tg/-)* were obtained by crossing *Pvalb-IRES-Cre* and *SOM-IRES-Cre* mice with the GP8.20 thy1-jRGECO1a line (Jackson Labs, JAX #030525). All lines were maintained in a C57Bl/6j background. All mice were fed *ad libitum* and housed on a 12-hour light/dark cycle in individual cages following craniotomies.

#### Viral injections

For interneuron silencing experiments, *PV-Cre(Cre/wt)::thy1-jRGECO1a(Tg/-)* and *SOM-Cre(Cre/wt)::thy1-jRGECO1a(Tg/-)* transgenic mice received an intracortical injection of 1.5×10^12^ vg/mL AAV2/8-hSyn-DIO-HA-hM4D(Gi)-IRES-mCitrine (Adgene, plasmid #50455) following skull removal (S1 : -2.0 mm AP; -3.2 mm ML from Bregma, 200 – 800 μm depth). Each mouse received a total dose of 1.2×10^9^ vg delivered in 800 nL : 200nL were injected every 200 μm from 800 μm to 200 μm to maximise transduction across cortical layers.

#### Chronic window implantation

Chronic cranial windows were implanted over the right barrel cortex (vS1) as previously described^1^. In brief, before the surgery, mice were injected with buprenorphine extended-release (subcutaneous (SC), 1.0 mg/kg). Animals were then anesthetized by ketamine–medetomidine injection (intraperitoneal (IP), 1 mg/kg and 0.4 mg/kg, respectively) and placed on a stereotaxic frame. An additional dose of ketamine (IP, 0.33 mg/kg) was administered as necessary to maintain anesthesia. Marcaine (SC,4 mg/kg) was injected locally under the scalp to provide local anesthesia. A titanium head bar was fixed to the skull over the cerebellum and occipital cortex using surgical glue (VetBond, CDMV, #188) and photo-curable dental cement (Flow-It ALC Flowable Composite, 1 ml syringe, catalog number N11VE, Patterson Dental). A craniotomy (about 3 mm × 3 mm) was performed over the right hemisphere using a dental drill. Room temperature 0.9% saline was regularly applied to prevent heating during the drilling process. A cover glass (0.17 mm thick) was placed over the cortex and sealed using dental cement. Throughout the surgery, medical O2 (1–2 L/min) was administered through a nose cone, and temperature was monitored using a rectal probe and maintained between 35 °C and 37 °C. After the surgery, mice were administered atipamezole (SC, 2 mg/kg), dexamethasone (SC, 2 mg/kg), meloxicam (SC, 2 mg/kg, for 2 days), enrofloxacin (SC, 5 mg/kg, every day for 3 days) and were allowed to recover for 3 weeks before experimentation.

#### Sedation protocol

For sedated imaging experiments, mice were first anesthetized using isoflurane (2% in 100% medical O2) in an induction box (SomnoSuite, Kent Scientific). Animals were then transferred to the head fixation apparatus, where isoflurane was administered through a nose cone. A bolus of dexmedetomidine (Dexdomitor, SC, 0.0925 mg/kg) was injected, followed by a continuous infusion of dexmedetomidine (0.185 mg/kg/hour) through a subdermal 24-gauge catheter (SURFLO Fluoropolymer Resin I.V. Catheters, Thermo Fisher Scientific, #22-251491) during the entire experiment. Isoflurane was gradually withdrawn by reducing the concentration by 0.5% every 10 min. Imaging began 10 min after isoflurane was reduced to 0% and the O2 nose cone removed. Temperature was monitored using a rectal probe and maintained between 35 °C and 37 °C during the entire imaging session using a feedback-controlled electric heating pad. Following the experiment, mice received atipamezole (SC, 2mg/kg) and cages were placed over a heating pad for recovery.

#### Whisker stimulation

All whiskers on the left side of the snout except for the number 2 column (A2–E2) were trimmed under 2 % isoflurane anesthesia prior to imaging. The mouse (awake or anesthetized) was then placed on a custom-made head fixation setup, and a single whisker was threaded through a glass capillary tube (OD: 1.2 mm, ID: 0.69 mm, length: 75 mm) attached to a bending piezoelectric element (Thorlabs, cat #PB4NB2W). The deflection was initiated by a piezo amplifier (Thorlabs, MDT693B), controlled by a custom-made microcontroller (Arduino, UNO Rev3) synchronized with the imaging software. Each whisker stimulation repetition consisted of a 4 second pre-stimulation period (baseline), a 4 second stimulation, a 10 second post-stimulation period and a 2 inter-trial window. For awake experiments, the post-stimulation period was reduced to 5 seconds. Each stimulation series consisted of 30 trials, randomly alternating between ‘real’ and ‘sham’. Each trial had a 30% probability of being a ‘sham’.. Stimulation trains consisted of 120 Hz cosine waveforms, repeated at 5 Hz. A 4-kHz tone (0.5 s) was delivered to signal the beginning of the reward availability window, even for sedated experiments. For sedated functional mapping experiments, E2, D2, C2 and B2 were sequentially stimulated. Only D2, C2 and B2 were stimulated for awake mapping experiments, as E2 could not be reliably threaded due to its angle on the mouse snout and the animal’s natural tendency to whisk during the threading when awake. In experiments where multiple whiskers were stimulated, the piezo was positioned next to the snout so that the capillary tube was placed vertically to allow for the simultaneous deflection of all whiskers between E2 and B2. For all silencing experiments, the stimulated whisker was selected *a priori,* based on the mCitrine expression within the associated barrel.

#### Interneuron silencing experiments

The hM4D(Gi) receptor was activated via intraperitoneal injection of the DREADD agonist deschloroclozapine (DCZ, 0.1 mg/kg)^30,31^. During syringe preparation, 2µL of DCZ stock (10 mg/mL) was diluted into 1mL of sterile saline, for a final concentration of 0.02 mg/mL. For all sedated experiments, a ‘baseline’ series was first recorded, followed by a second series 20 minutes post-DCZ administration. For all awake experiments, the post-DCZ recording was compared to a post-vehicle injection recording, acting as control (saline).

#### Wide-field imaging and analysis

Wide-field imaging and analysis was performed using previously described methods^1^. Briefly, intrinsic optical signals and fluorescence images were acquired with an optical imaging system (OiS200, LabeoTech) consisting of a 1024 × 1024 sCMOS camera, a camera lens (50-mm f/1.2) and filtered LEDs. For intrinsic signals, the brain was illuminated using a green (centered at 525 nm) and a red (centered at 625 nm) LED. GCaMP6f fluorescence was excited using a band-pass filtered blue LED (centered at 475 nm, Semrock filter FF02-472-SP), whereas jRGECO1a fluorescence was excited using a band-pass filtered lime LED (centered at 550 nm, Semrock filter FF01-554/23). Backscatter and fluorescence signals were filtered through a multi-band-pass filter (FF01-512/630, Semrock) placed in front of the camera. Data were collected using LabeoTech’s IOI imaging software. Camera frames were acquired at 60 Hz, with a 512 × 512 resolution (2 × 2 binning), while LEDs were strobed, for a final frame rate of 20 Hz per channel. Exposure time was 1 ms for backscatter signals and 6 ms for fluorescence. For each experiment, LED power was adjusted to minimize undersaturation and oversaturation for each channel and was kept constant throughout the experiment. Image focus was adjusted by moving the entire apparatus (camera and lens) with a manual linear stage. At the beginning of each experiment, a snapshot of the surface vasculature was taken for alignment purposes. Then, the focus was adjusted to approximately 400 µm below the surface for imaging. Image analysis was performed with custom-made MATLAB scripts (MathWorks), and scripts were adapted from the UMIT MATLAB suite (LabeoTech).

During signal processing, each channel was first de-trended using previously described methods (double exponential equation)^1^. Fluorescence signals were then corrected for changes in hemoglobin absorption (hemodynamic correction) using the pixelwise regression method based on the two backscatter wavelengths (525 nm and 625 nm)^43^. Pixels were then normalized to a baseline of 3 seconds before stimulation, and a Gaussian filter was applied across space (σ = 3). Changes in HbO, HbR and HbT concentrations were estimated using the modified Beer–Lambert law as previously described^43,44,45^. Averages were made from a minimum of 15 trials. ROIs, roughly corresponding to the cortical representation of each whisker, were obtained by thresholding (85% threshold) the average GCaMP6f or jRGECO1a response between stimulation onset (t = 4 seconds) and 2 seconds after stimulation (t = 10 seconds). Standardized ROIs were drawn based on the Allen Institute Brain Atlas, as previously described^46^, and were registered onto the centroid of the Ca^2+^ signal corresponding to each whisker-stimulation.

Ca^2+^ and hemodynamic signals were smoothed using a 250 ms and 500 ms moving average respectively. Responses to single-whisker stimulation were extracted from each ROI and expressed as a function of their position along the 2nd barrel column relative to the stimulated barrel (associated barrel, 1 barrel away, 2 barrels away, 3 barrels away). Responses to multiple-whisker stimulation were extracted from an ROI encompassing the 4 stimulated barrels, E2-B2.

The overall area under the curve was calculated within a 4 second time interval for Ca^2+^ signals (t = 4 seconds - t = 8 seconds) and within a 6 second time interval for hemodynamic signals (t = 4 seconds - t = 10 seconds). When analyzing the early phase of the response, signals were considered within a 0.5 seconds time window for calcium signals (t = 4 seconds - t = 4.5 seconds) and a 1 second time window for hemodynamic signals (t = 5.3 seconds – t = 6.3 seconds). Analysis window was shifted 1.3 seconds for hemodynamic signals based on a hemodynamic response function (HRF) model prediction^27,47^.

Spatial spread analysis was achieved by gradually expanding donut-shaped masks of 25 µm radius around an initial ROI centered around the peak of the jRGECO1a activity. The response AUC was calculated within each circular mask and normalized on a scale of 0 to 1, with 1 being assigned to the donut that displayed the maximal response and 0 being assigned to the donut that displayed the smallest response. Signals were standardized over 40 donuts for interneuron Ca^2+^ signals analysis (Fig. 2), while 120 were used to analyze the spread of excitatory Ca^2+^ and hemodynamic signals in chemogenetic experiments as the hemodynamic response spread further away from the stimulated barrel following PV silencing (Fig. 5 and Supplementary Fig. 5). The half-max radius consisting of the distance under which the signal goes below a threshold of 50% of the maximum value was calculated. The primary visual cortex (V1) and all secondary (extrastriate) visual areas were excluded from this analysis.

#### Two-photon microscopy and data analysis

Prior to imaging, jRGECO1a-expressing mice were retro-orbitally injected under 2% isoflurane anesthesia with either Alexa 680 NHS ester conjugated to 2,000 kDa amino dextran (100 µl at 3.5–5%, A20008, Thermo Fisher Scientific; AD2000x100 or AD2000x250, Fina Biosolutions) to label the vasculature. Two-photon microscopy was performed using a tilting Bergamo II system (Thorlabs) controlled through the ThorImage software and equipped with a dual-output, tunable femtosecond laser (Discovery NX-TPC, Coherent). The laser beam was focused through a 25X, 0.95 NA water immersion objective (Leica HC FLUOTAR L ×25/0.95 W VISIR) and power was attenuated using a built-in AOM and was kept under 100 mW to prevent tissue damage. Laser was scanned horizontally using an 8-kHz galvo-resonant (1024 x 512 pixels, bi-directional scanning for a linear of 60Hz). Photons were detected with GaAsP photomultiplier tubes (PMT2100, Thorlabs).

Viral expression spread was estimated by capturing *z-stacks* of layer 2/3 DREADD-positive interneurons (mCitrine, excited at λ=925nm) along with *z-stacks* of the vasculature (Alexa 680, excited at λ=1280nm). Alexa 680 and mCitrine emissions were split and band-pass filtered using a 562 nm dichroic mirror (FF562-Di03, Semrock) and a set of emission filters (FF01-685/LP-25 and FF03-525/50 respectively, Semrock). To map hM4Di-expressing interneurons onto the barrel map obtained during wide-field imaging, corresponding control points were selected on the two-photon maximum intensity projection of the surface vasculature and the wide-field reference image of the cranial window (reflectance at 525 nm) for co-registration. The x–y position of mCitrine-positive interneurons was then marked on a z-stack of the imaged region and transposed to the wide-field barrel map using a transformation vector. To measure spontaneous calcium transients in layer 2/3 pyramidal cells, jRGECO1a was excited at 1125 nm. Alexa 680 and jRGECO1a emissions were split by a 660 nm dichroic mirror (FF-660-Di02, Semrock) and band-pass filtered using a set of emission filters (FF01-685/LP-25 and FF02-617/73 respectively, Semrock). Cell bodies and neuropil were segmented using the *Suite2p* software^48^. Automatic ROI masks detection was used, but additional masks could be added after verification. A neuropil donut was automatically traced around each soma with an inner radius of two pixels and a minimum total of 350 pixels. No overlap between somas was tolerated during the segmentation process. A coefficient of 0.7 was used for neuropil subtraction. Calcium transients were calculated as Δ*F/F* = (*F − F_0_*) / *F_0_*, where *F_0_* represents baseline fluorescence and *F* the fluorescence at time *t*. The standard deviation was calculated over the first recording (baseline series, pre-DCZ injection). All peaks above a 3-standard deviation threshold from a 100ms local baseline window were considered as an individual calcium transient.

#### Statistics

Statistical tests were performed using the GraphPad Prism 10 software. When analyzing the effect of one or more independent variables, multiple linear regressions (MLR) were used. The goodness of the fit was measured using the adjusted R^2^ *(R^2^).* The statistical tests used for each experiment and relevant *p* values are represented on the figures and/or are reported in the text or figure legends, where appropriate. For comparing data across two or more groups, a two-tailed Student’s t-test or a one-way ANOVA followed by a Tukey’s multiple comparison post hoc test was used. The effect of two independent, categorical variables was analyzed using a two-way ANOVA or a mixed-effect model when some data points were missing followed by Tukey’s multiple comparison or Sidak’s multiple comparisons post hoc test. Full results for statistical tests and MLR models, including exact *p* values and the list of variables included in each MLR model, are available in Supplementary Information. Sex and age were not considered in any statistical test or analysis. All data shown in the manuscript are represented as mean ± SEM.

## Acknowledgements

The authors thank Dr. Richard Robitaille and Dr. Jean-Claude Lacaille for supplying the PV-Cre and SOM-Cre mice, as well as Pierette Kwemo and Isabel Laplante for colony management. RLR holds a Canada Research Chair in Neurovascular Interactions (RLR). The work was funded via a Natural Sciences and Engineering Research Council of Canada discovery grant RGPIN-2020-05276 (RLR) and a Brain Canada Momentum Grant to RLR in partnership with the Heart and Stroke Foundation of Canada. RLR was also funded from CIHR project grants No. 451469 and 519817. LZ had a doctoral scholarship from CIRCA, AM holds a FRQS doctoral training award, and MRL holds a CIHR postdoctoral fellowship.

## SUPPLEMENTARY FIGURES

**Supplementary Figure 1.**
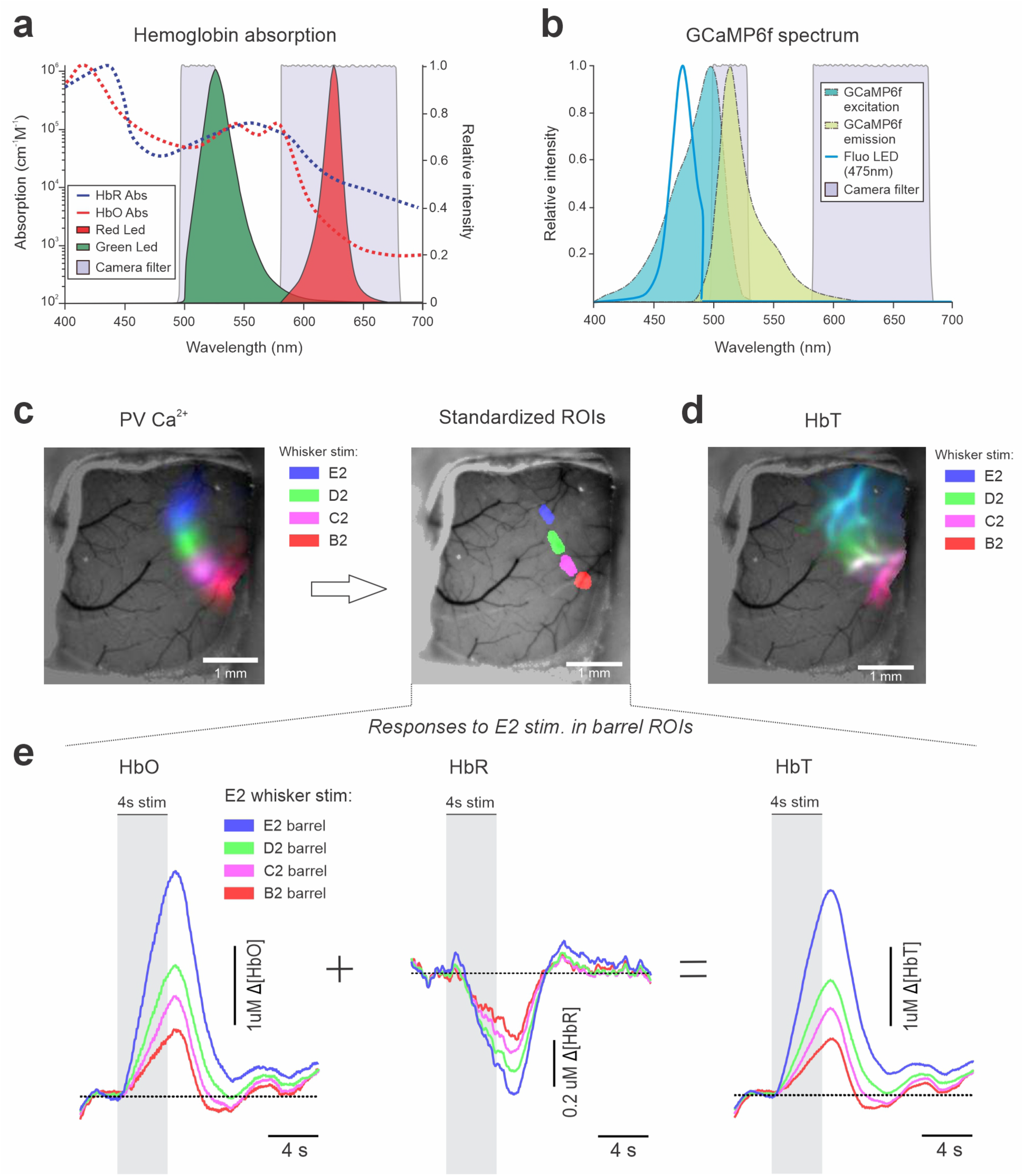
Mesoscale imaging of interneuron Ca^2+^ and hemoglobin dynamics. **a,** Absorption spectra of HbO and HbR in dotted lines, as well as the spectra of the green and red LEDs including their filters used to estimate changes in hemoglobin concentration. **b,** GCaMP6f excitation and emission spectra as well as the spectrum of the filtered blue LED, centered at ∼475nm, used to excite GCaMP6f in interneurons. **c,** Example images of average changes GCaMP6f (left) fluorescence in response to the stimulation of four individual whiskers (color-coded) in a PV-GCaMP6f mouse, same as in Figure 1. Standardized barrels ROIs (right) were aligned to centroids of the thresholded (85%) GCaMP6f signal. **d,** Example image of average changes in HbT in response to the stimulation of four individual whiskers (color-coded). **e,** Example of HbO, HbR and HbT traces in different barrel ROIs in response to the stimulation of whisker E2 in the same mouse shown in (**c**) and (**d**).

**Supplementary Figure 2.**
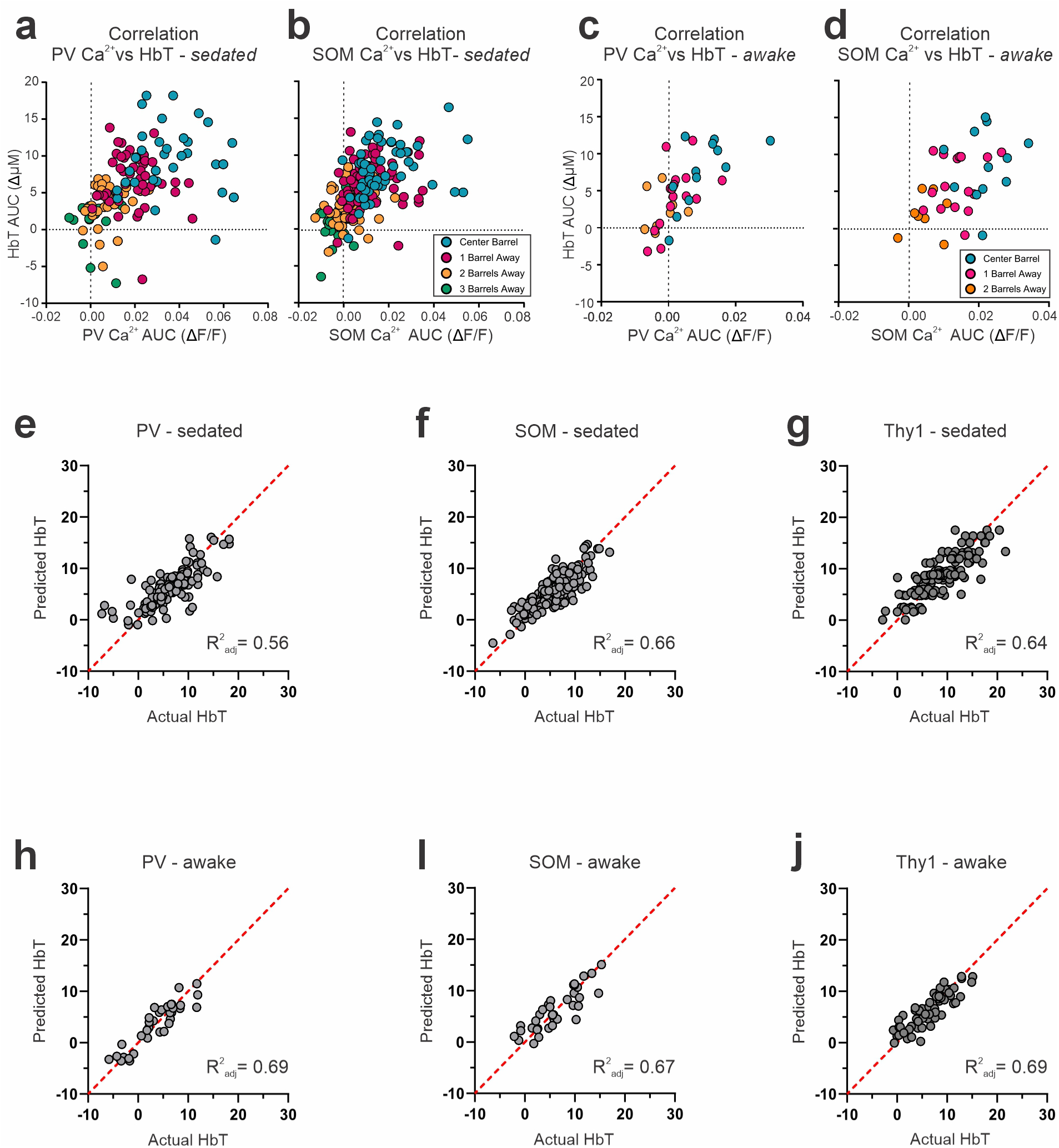
HbT variance is explained by interneuron calcium signals. **a-d,** Changes (AUC) in Ca2+ (ΔF/F) and HbT (µM) per barrel ROI for sedated *PV-GCaMP6f* (**a**) and *SOM-GCaMP6f* (**b**), as well as awake *PV-GCaMP6f* (**c**) and *SOM-GCaMP6f* (**d**). **e-j,** Portion of HbT variance that can be predicted by Ca^2+^ in sedated PV (**e**), sedated SOM (**f**), sedated Thy1 (**g**), awake PV (**h**), awake SOM (**i**), awake Thy1 (**j**) versus actual HbT measurements. **a-j,** For AUC changes, AUC _t=4-8s_ was used for Ca^2+^ signals and AUC _t=4-10s_ was used for HbT. *PV-GCaMP6f* - sedated: N=9, 3M and 6F; awake: N=4, 3M and 1F. *SOM-GCaMP6f -*sedated: N=12, 4M and 8F; awake: N=4, 2M and 2F.

**Supplementary Figure 3.**
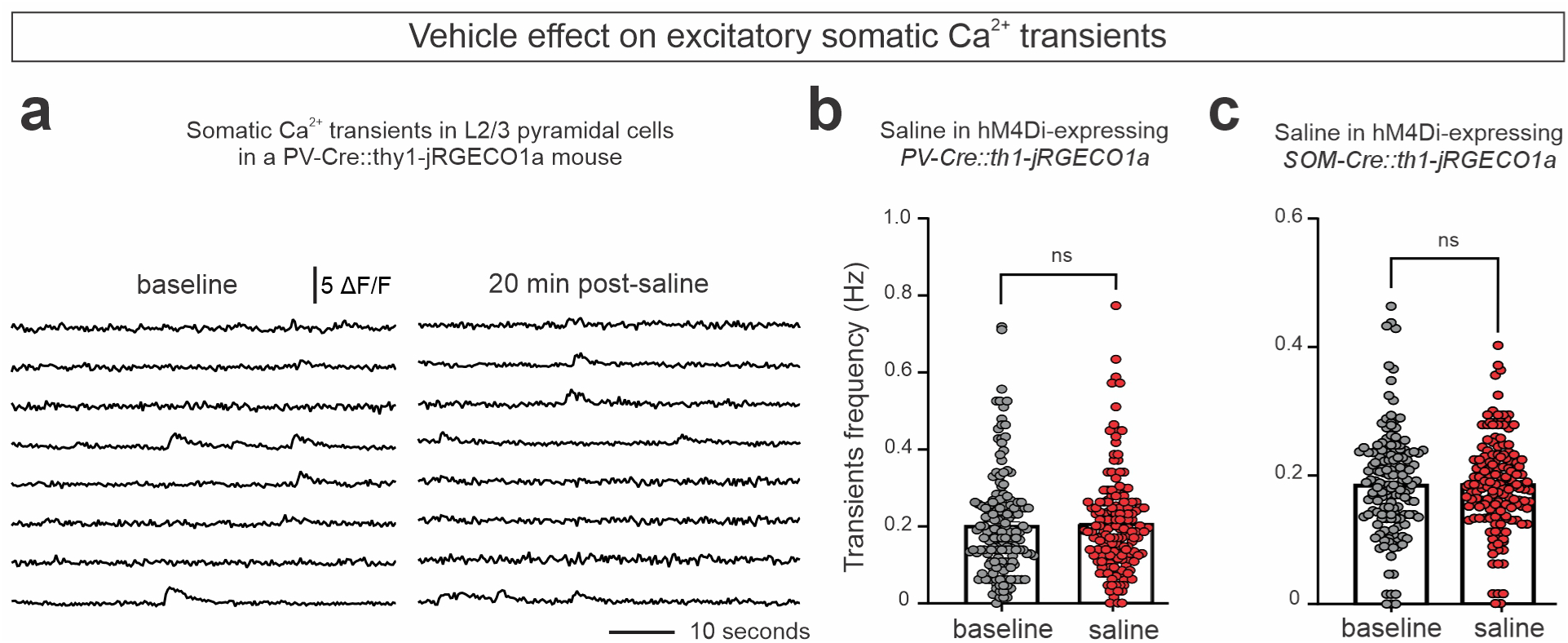
Saline (vehicle) injections do not affect excitatory Ca^2+^ transient frequency. **a,** Example of spontaneous Ca^2+^ transients measured in the somas of eight L2/3 pyramidal cells before (left) and 20 minutes post-saline injection (right) in an hM4Di-expressing *PV-Cre::thy1-jRGECO1a* mouse, measured in ΔF/F. **b,c,** Ca^2+^ transient frequency in Hz before and after saline injection in hM4Di-expressing *PV-Cre::thy1-jRGECO1a* (**b**, N=3, 2M and 1F) and *SOM-Cre::thy1-jRGECO1a* mice (**c**, N=5, 5M). ns: *p*>0.05 by two-tailed pared t-test. Between 58 and 168 somas per mouse were included.

**Supplementary Figure 4.**
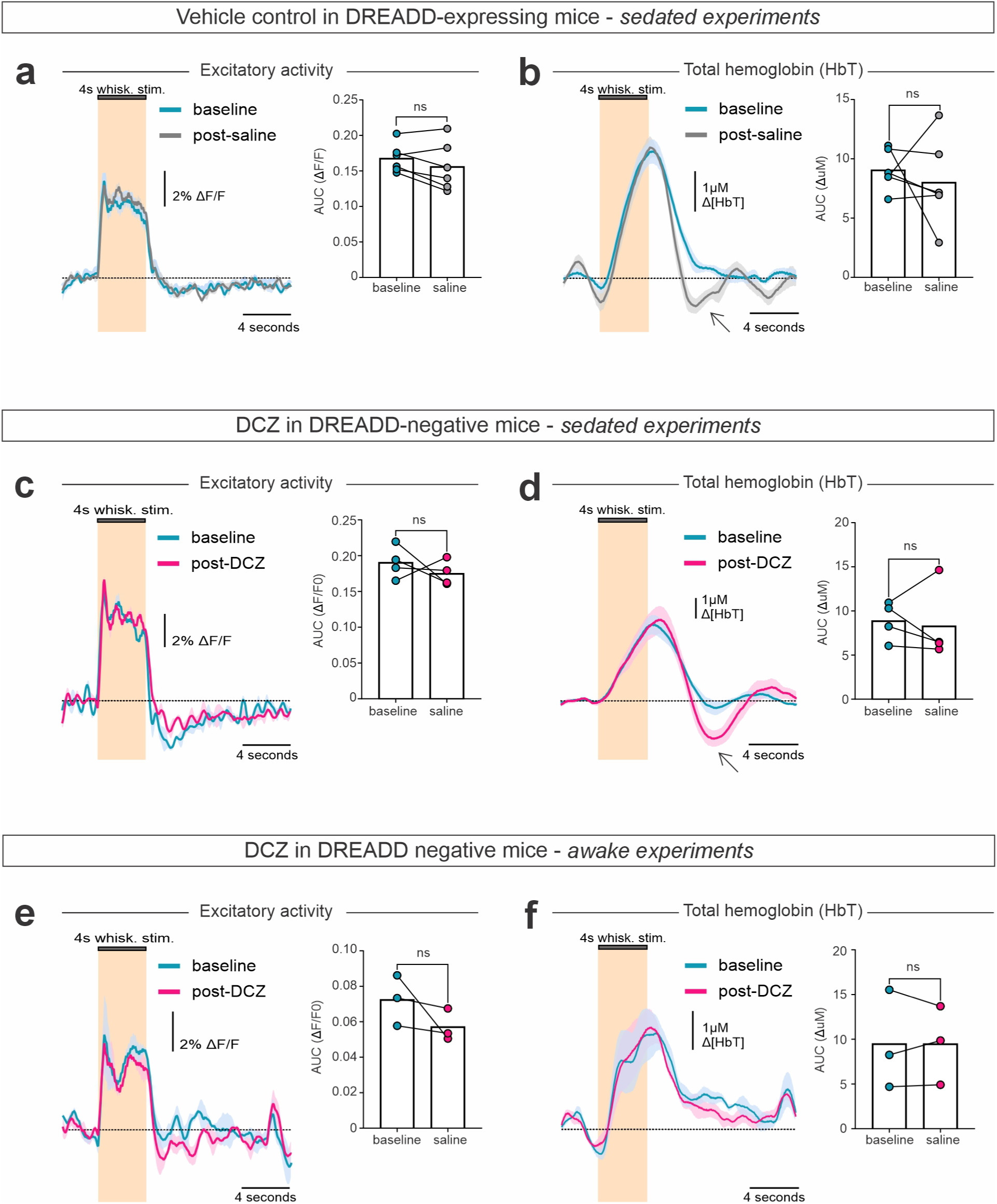
Vehicle and DCZ controls on NVC. **a,b,** Traces (left) and AUC (right) of average excitatory Ca^2+^ (**a**) and hemodynamic (**b**) responses to a single whisker stimulation before and after vehicle injection in sedated hM4Di-expressing *PV-Cre::thy1jRGECO1a mice* (N=6, 5M and 1F). **c,d,** Traces (left) and AUC (right) of average excitatory Ca2+ (**c**) and hemodynamic (**d**) responses to a single whisker stimulation before and after DCZ injection in sedated hM4Di-negative *PV-Cre::thy1jRGECO1a* mice (N=4). **e,f,** Traces (left) and AUC (right) of average excitatory Ca2+ (**e**) and hemodynamic (**f**) responses to a single whisker stimulation before and after DCZ injection in awake hM4Di-negative *PV-Cre::thy1jRGECO1a* mice (N=3). **a-f,** Data are represented as the mean ± the SEM. Shaded areas on signal traces represent the SEM. ns: *p*>0.05 by two-tailed paired t-test. **b,d**, Arrow points to change in post-stimulus undershoot following both saline and DCZ in sedated mice.

**Supplementary Figure 5.**
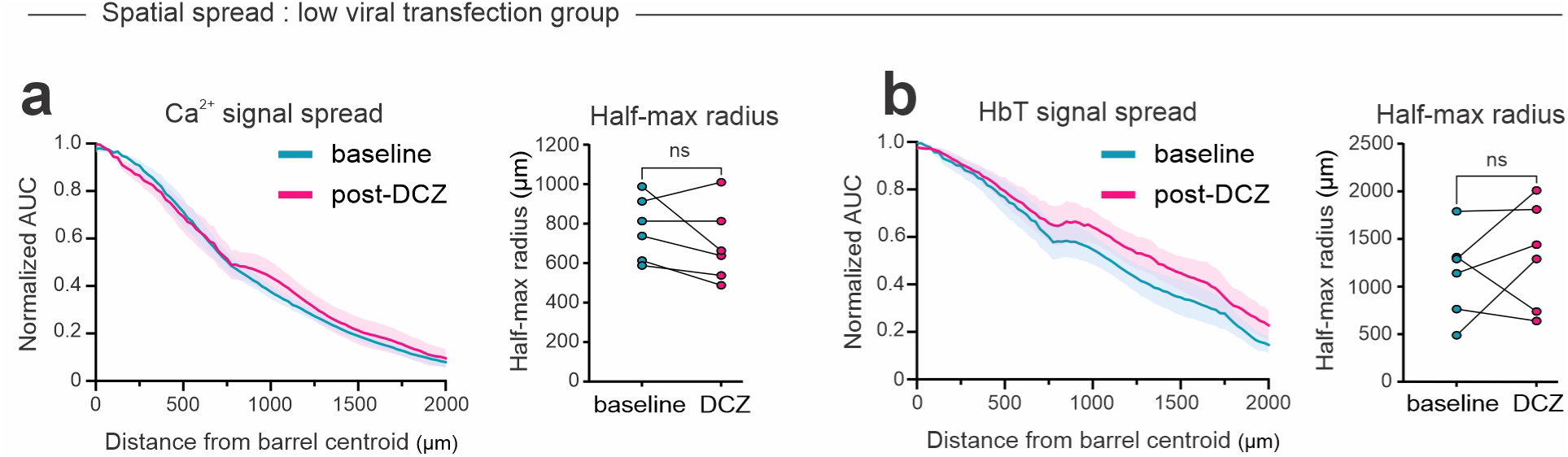
Chemogenetic PV silencing does not alter the spatial dynamics of NVC in the low spread viral expression group. **a,b,** Excitatory Ca^2+^ and HbT signal spread (Normalized AUC) from the centroid barrel and half-max radius (μm) in low spread hM4Di-expressing *PV-Cre::thy1-jRGECO1a* mice. All analysis was done on the early phase Ca^2+^ AUC _t=4-4.5s_ and hemodynamic response AUC _t=5.3-6.3s_. N=6, 2M and 4F

**Supplementary Figure 6.**
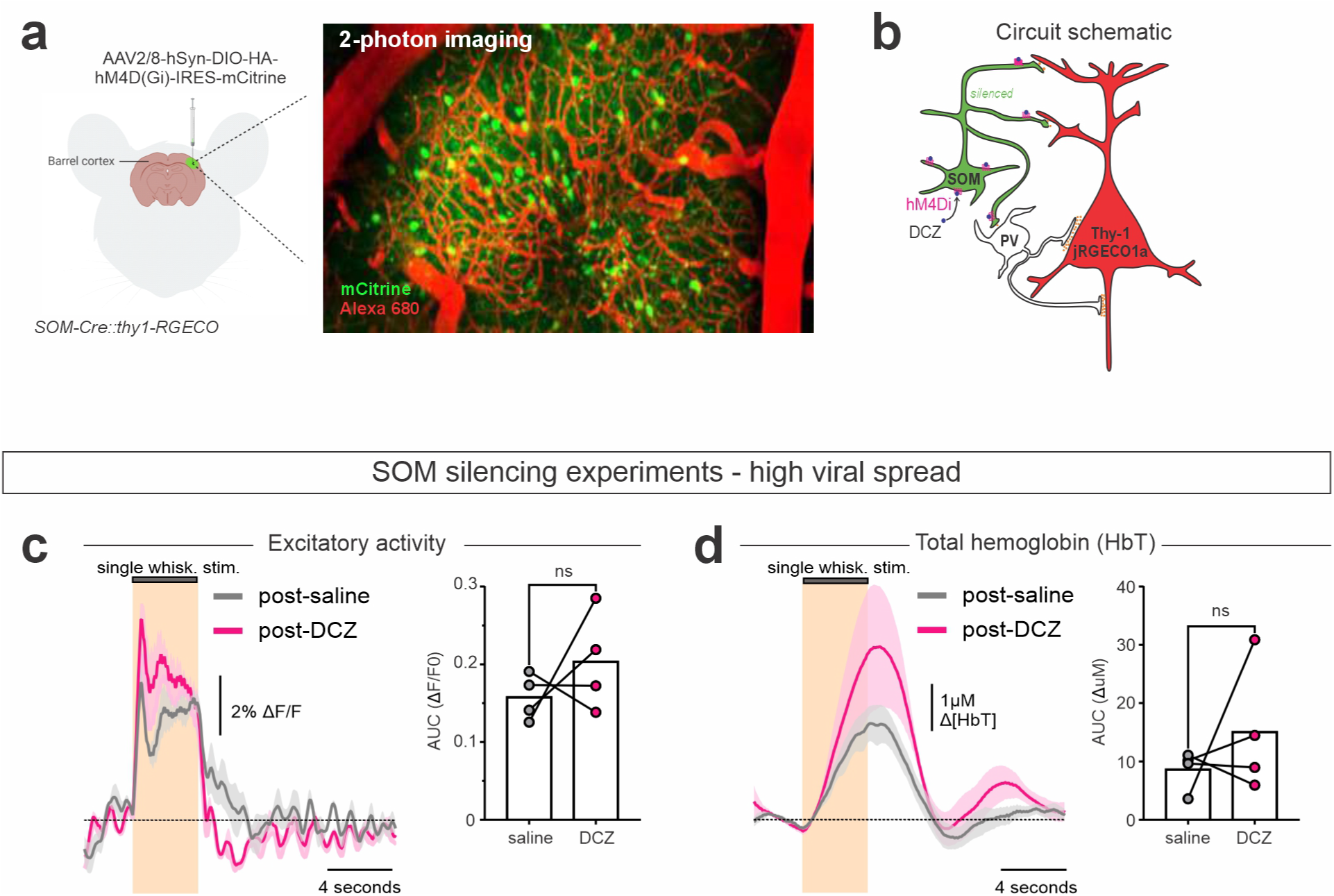
Supplementary data from the high spread viral expressing group in SOM interneurons. **a,** SOM silencing experiment schematic. The Cre-dependent AAV2/8-hSyn-DIO-HA-hM4D(Gi)-IRES-mCitrine was locally injected in S1 of *SOM-Cre::thy1-jRGECO1a* mice. Viral expression was confirmed via localization of mCitrine-positive interneurons using two-photon microscopy. Blood vessels were labeled with intravenous Alexa680-dextran. **b,** Schematic of the circuit. SOM interneurons, co-expressing the hM4D(Gi) receptor and mCitrine (green). To selectively silence SOM interneurons, DCZ is injected to activate the hM4D(Gi) receptor while recording the impact on excitatory neuron Ca2+, via jRGECO1a fluorescence (red), as shown in Fig. 3. **c,d,** Traces (left) and AUCs (right) of average excitatory Ca^2+^ (**c**) and hemodynamic (**d**) responses to a single whisker stimulation after saline (vehicle) or DCZ injection in sedated *SOM-Cre::thy1jRGECO1a* mice with high spread viral expression (i.e. ≥ 2 barrels, N=4/6, 3M and 1F). Data are represented as the mean ± the SEM. Shaded areas on signal traces represent the SEM. ns: *p*>0.05 by two-tailed paired t-test.

